## Supplementary Figures 1 - 12 for "Structural and functional asymmetry of the neonatal cerebral cortex"

### This PDF file includes:

Figs. S1 to S12
SI References

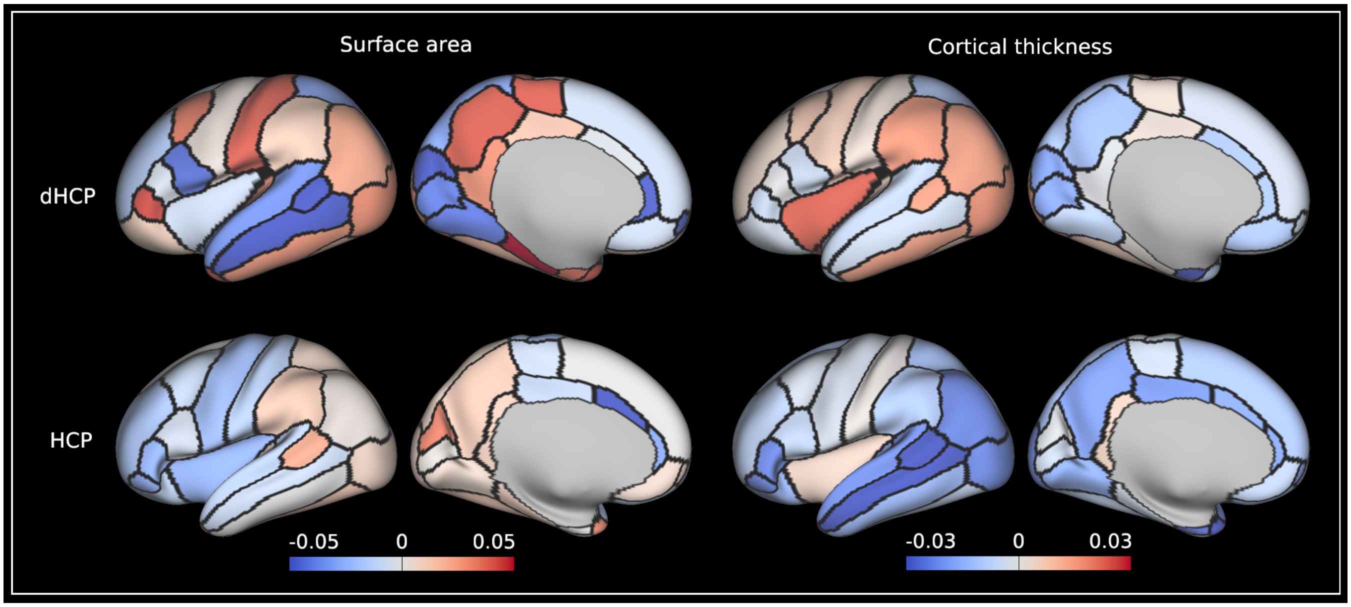


**Fig. S1.** Asymmetry indices of surface area and cortical thickness for the dHCP and HCP-YA cohorts, averaged (for the dHCP) within regions of the the M-CRIB-S atlas (neonatal equivalent of Desikan-Killiany atlas) (1, 2) and (for HCP-YA) the Desikan-Killiany atlas (3). Leftward asymmetry indices are color-coded red, and rightward asymmetry indices are color-coded blue. Data at https://balsa.wustl.edu/1pn1V.

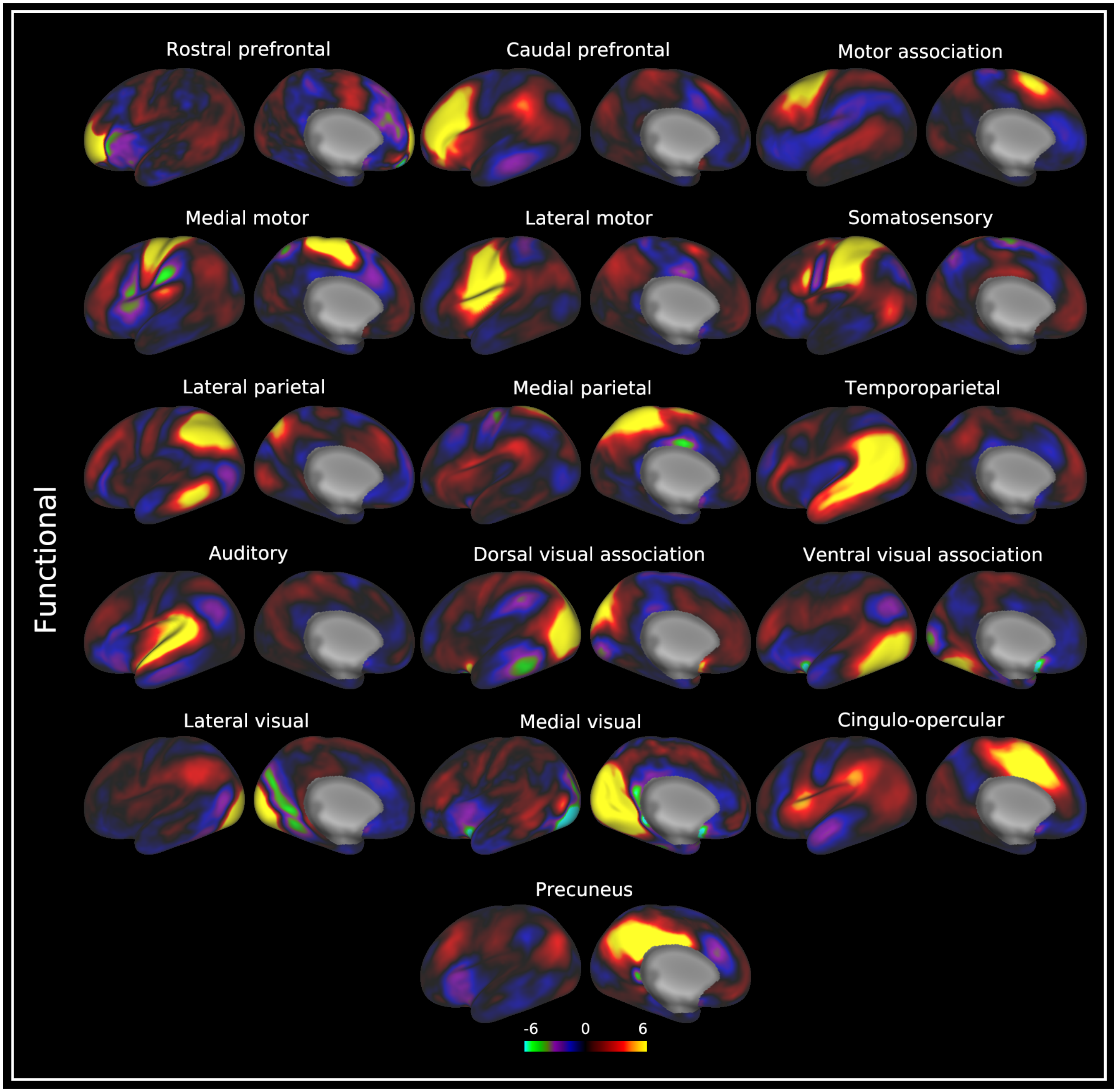


**Fig. S2.** Symmetric group resting-state networks for the dHCP. Resting-state network spatial maps visualised on very inflated 40-week PMA left hemispheric surface. Only the left hemisphere is shown for illustrative purposes. Data at https://balsa.wustl.edu/5B3M3.

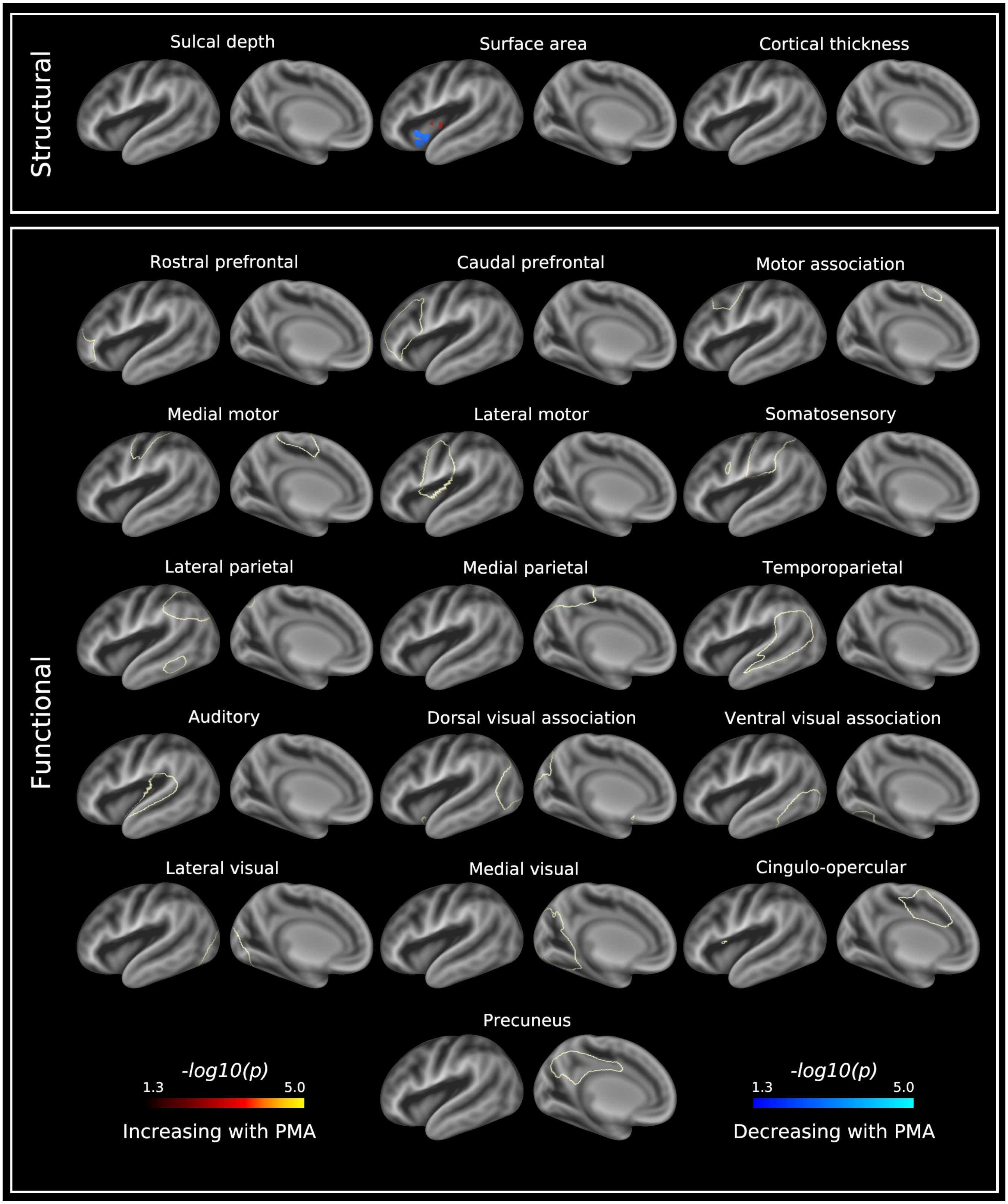


**Fig. S3.** $-\log_{10}(p)$-value maps for the effect of postmenstrual age on the structural and functional asymmetries in the healthy term-born neonatal cortex at term-equivalent age. Leftward asymmetries are represented by the red-yellow colour scale and rightward asymmetries by the blue-light blue colour scale. Significantly asymmetric regions are visualised on a very inflated 40-week PMA left hemispheric surface, and are overlaid on a 40-week PMA sulcal depth template (grey scale colour scheme). Off-white lines surrounding the functional asymmetries represent the mask used to threshold single subject asymmetry maps (see Methods: Generating Asymmetry Maps). Data at https://balsa.wustl.edu/n8Gvj.

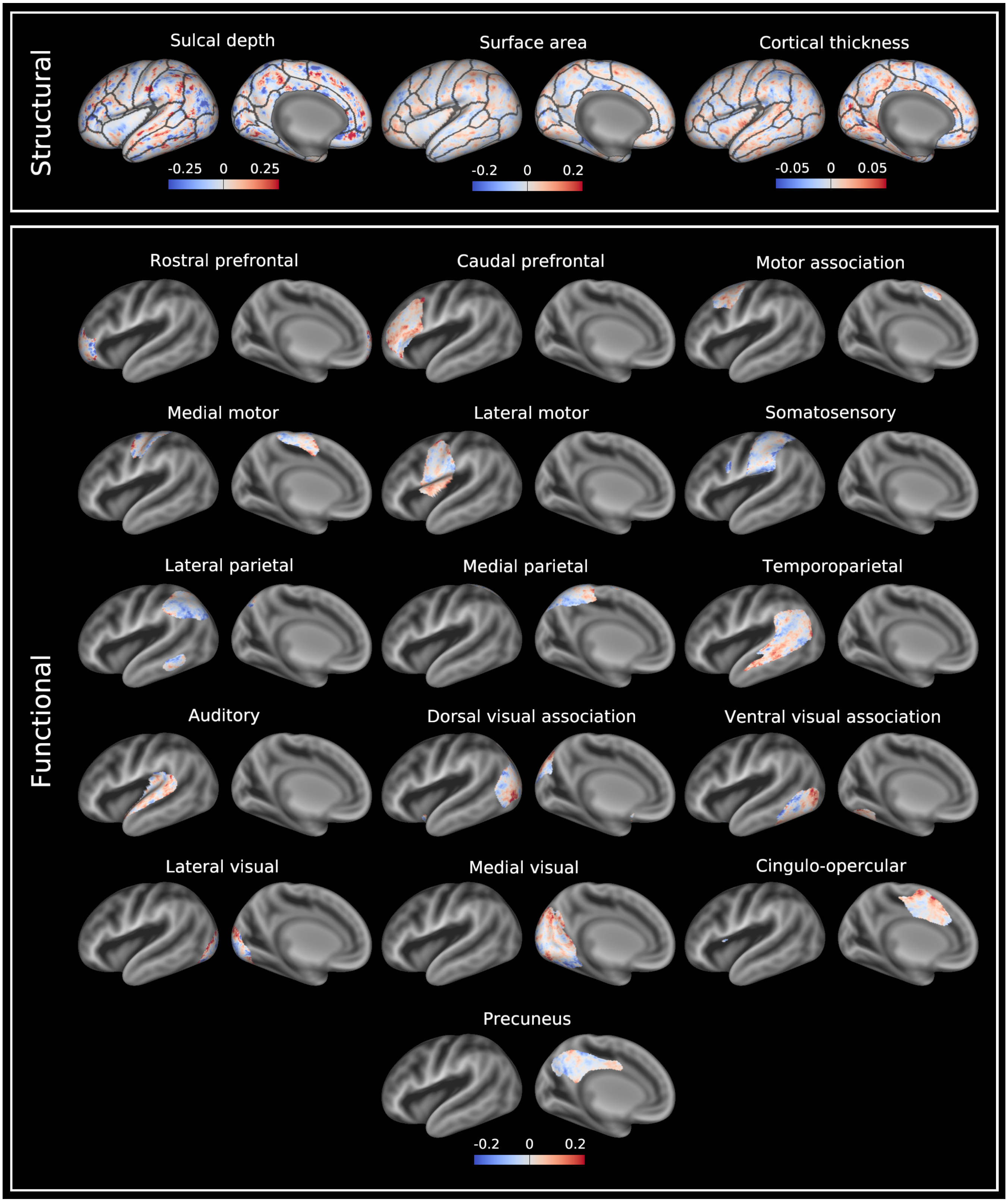


**Fig. S4.** Difference in median asymmetry indices (female - male) of structural and functional asymmetries across the healthy term-born neonatal cohort scanned at term-equivalent age. Leftward asymmetries are color-coded red, and rightward asymmetries color-coded blue. Asymmetry indices are visualised on a very inflated 40-week PMA left hemispheric surface, and are overlaid on a 40-week PMA sulcal depth template (grey scale colour scheme). Anatomical regions of interest from a neonatal version of the Desikan-Killiany atlas (M-CRIB-S) (1, 2) are overlaid on the structural asymmetries for reference.

Data at https://balsa.wustl.edu/gmgnv.

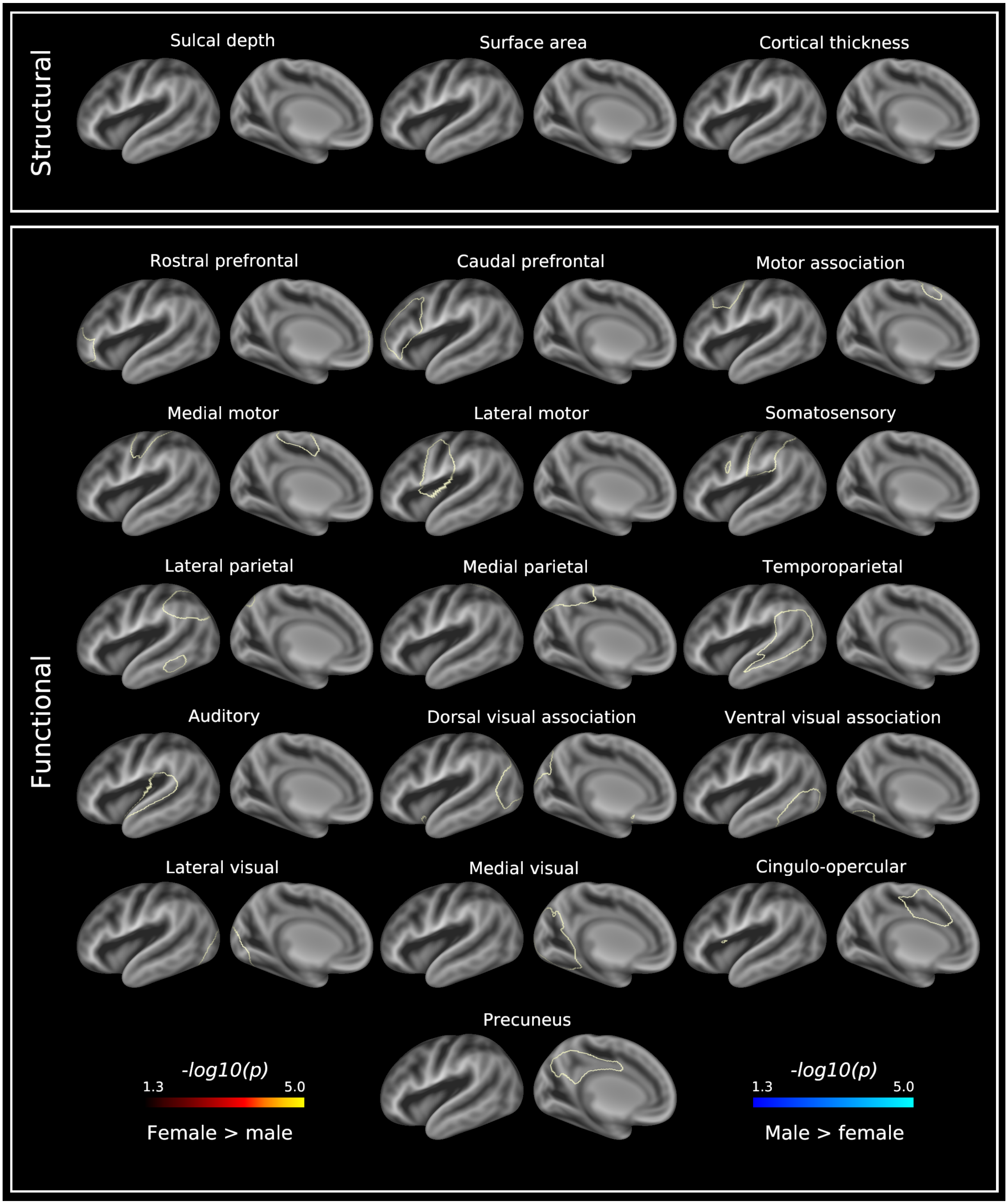


**Fig. S5.** $-\log_{10}(p)$-value maps for the effect of biological sex on the structural and functional asymmetries in the healthy term-born neonatal cortex at term-equivalent age. Leftward asymmetries are represented by the red-yellow colour scale and rightward asymmetries by the blue-light blue colour scale. Significantly asymmetric regions are visualised on a very inflated 40-week PMA left hemispheric surface, and are overlaid on a 40-week PMA sulcal depth template (grey scale colour scheme). Off-white lines surrounding the functional asymmetries represent the mask used to threshold single subject asymmetry maps (see Methods: Generating Asymmetry Maps). Data at https://balsa.wustl.edu/M9jX2.

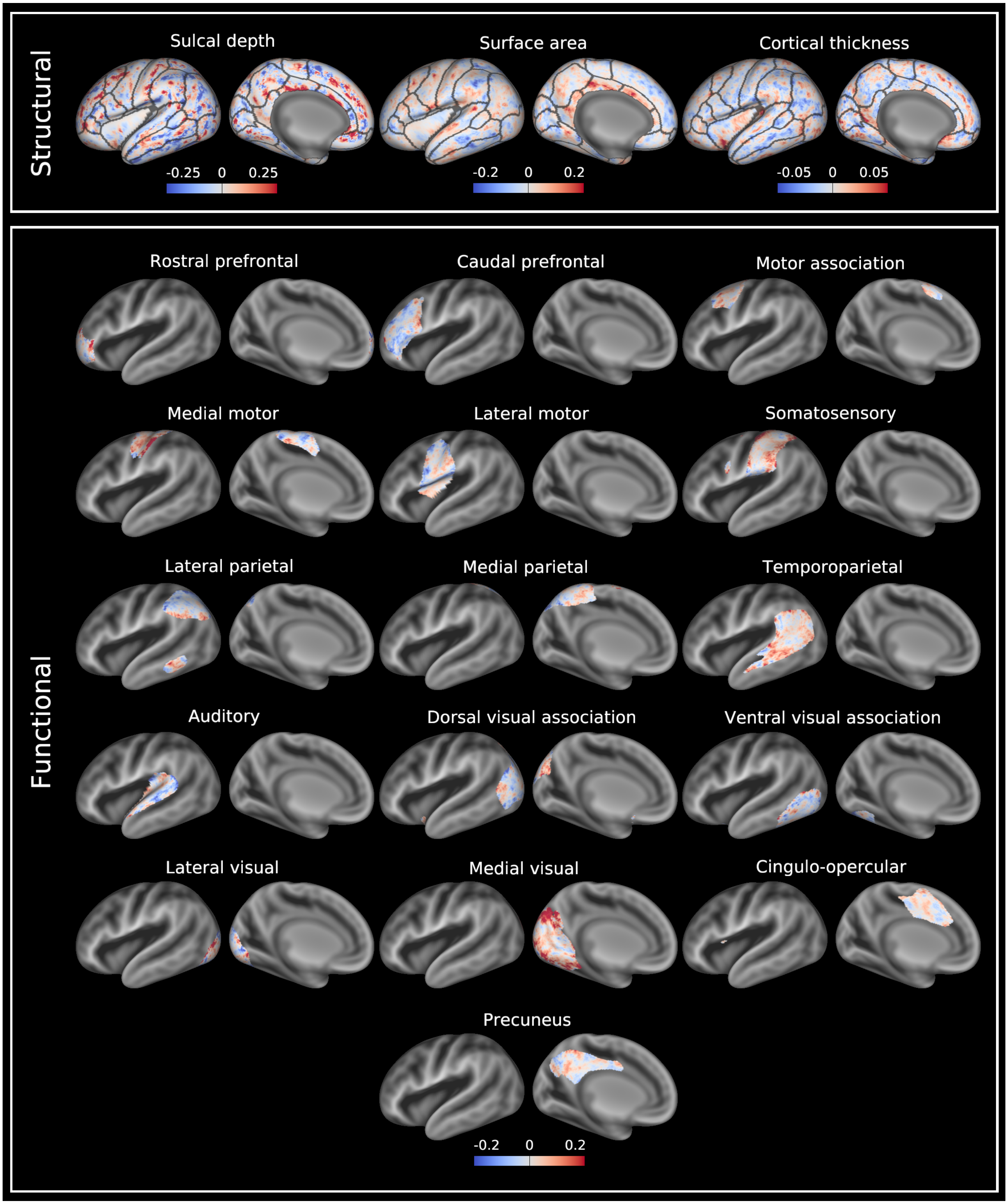


**Fig. S6.** Difference in median asymmetry indices of structural and functional asymmetries between term-born and preterm-born neonates (term - preterm) scanned at term-equivalent age. Leftward asymmetries are color-coded red, and rightward asymmetries color-coded blue. Asymmetry indices are visualised on a very inflated 40-week PMA left hemispheric surface, and are overlaid on a 40-week PMA sulcal depth template (grey scale colour scheme). Anatomical regions of interest from a neonatal version of the Desikan-Killiany atlas (M-CRIB-S) (1, 2) are overlaid on the structural asymmetries for reference.

Data at https://balsa.wustl.edu/Bg8Xm.

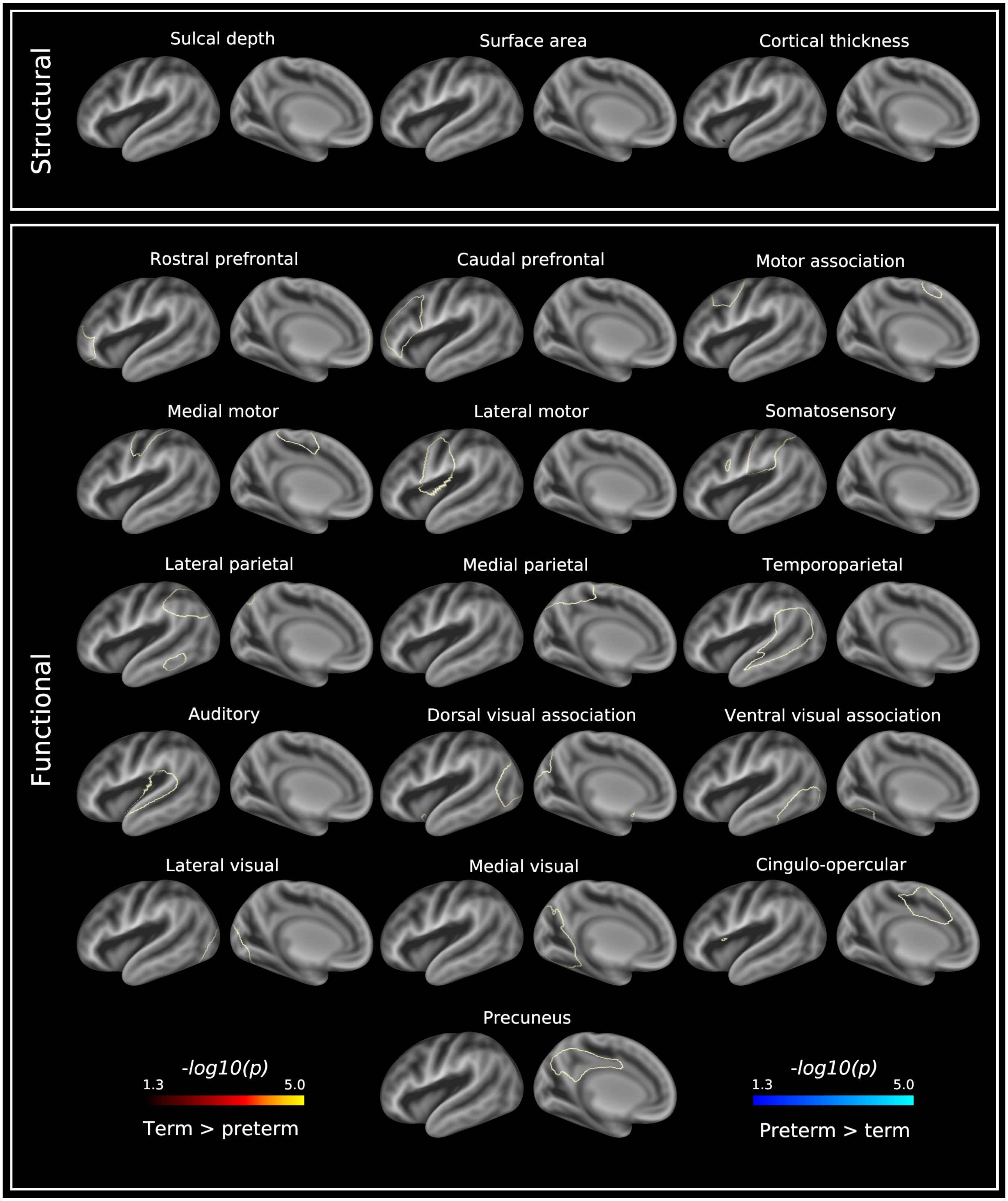


**Fig. S7.** $-\log_{10}(p)$-value maps for the effect of preterm birth on the structural and functional asymmetries in the neonatal cortex at term-equivalent age. Leftward asymmetries are represented by the red-yellow colour scale and rightward asymmetries by the blue-light blue colour scale. Significantly asymmetric regions are visualised on a very inflated 40-week PMA left hemispheric surface, and are overlaid on a 40-week PMA sulcal depth template (grey scale colour scheme). Off-white lines surrounding the functional asymmetries represent the mask used to threshold single subject asymmetry maps (see Methods: Generating Asymmetry Maps). Data at https://balsa.wustl.edu/l7gq0.

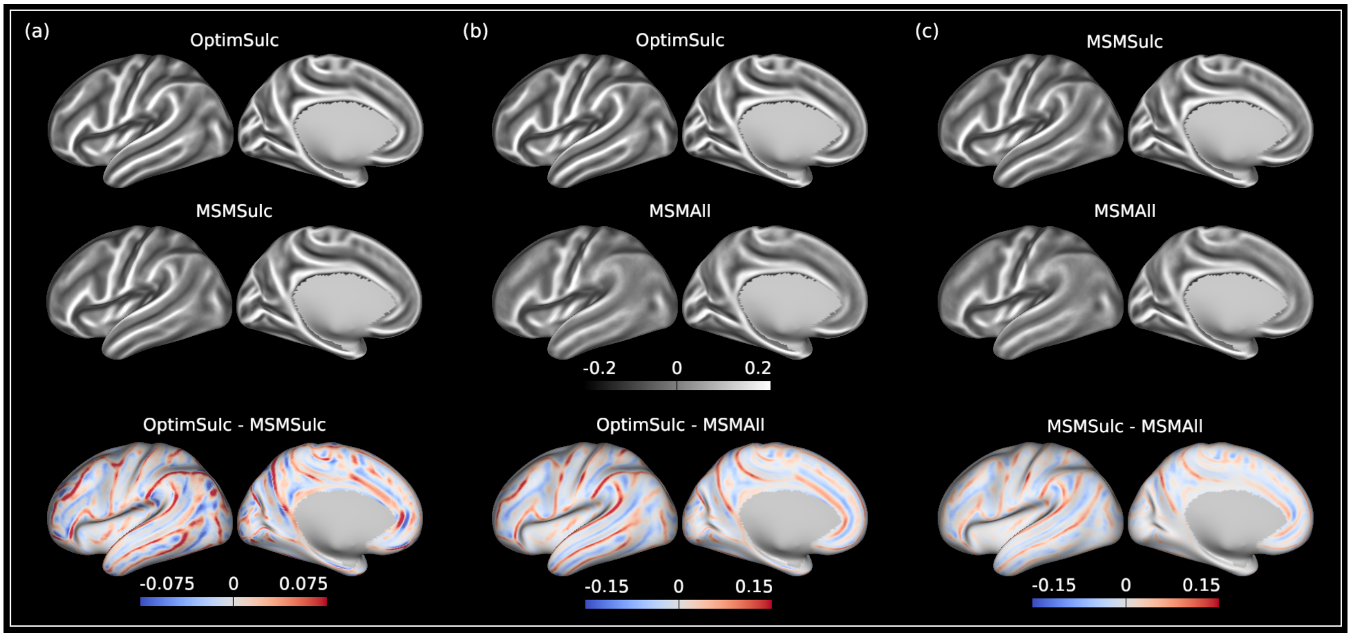


**Fig. S8.** Effect of surface registration on intersubject alignment of cortical folding. (a) Top row: median curvature maps across 1110 HCP-YA subjects after registration optimised for sulcal depth (OptimSulc); middle row: median curvature maps across 1110 HCP-YA subjects after MSMSulc registration; bottom row: difference in median curvature maps after OptimSulc and MSMSulc registration (OptimSulc - MSMSulc). (b) Top row: median curvature maps across 1110 HCP-YA subjects after OptimSulc registration; middle row: median curvature maps across 1096 HCP-YA subjects after MSMAll registration; bottom row: difference in median curvature maps after OptimSulc and MSMAll registration (OptimSulc - MSMAll). (c) Top row: median curvature maps across 1110 HCP-YA subjects after MSMSulc registration; middle row: median curvature maps across 1096 HCP-YA subjects after MSMAll registration; bottom row: difference in median curvature maps after MSMSulc and MSMAll registration (MSMSulc - MSMAll). Median curvature maps are visualised on very inflated left hemispheric surfaces for illustrative purposes. Data at https://balsa.wustl.edu/Bg0jv.

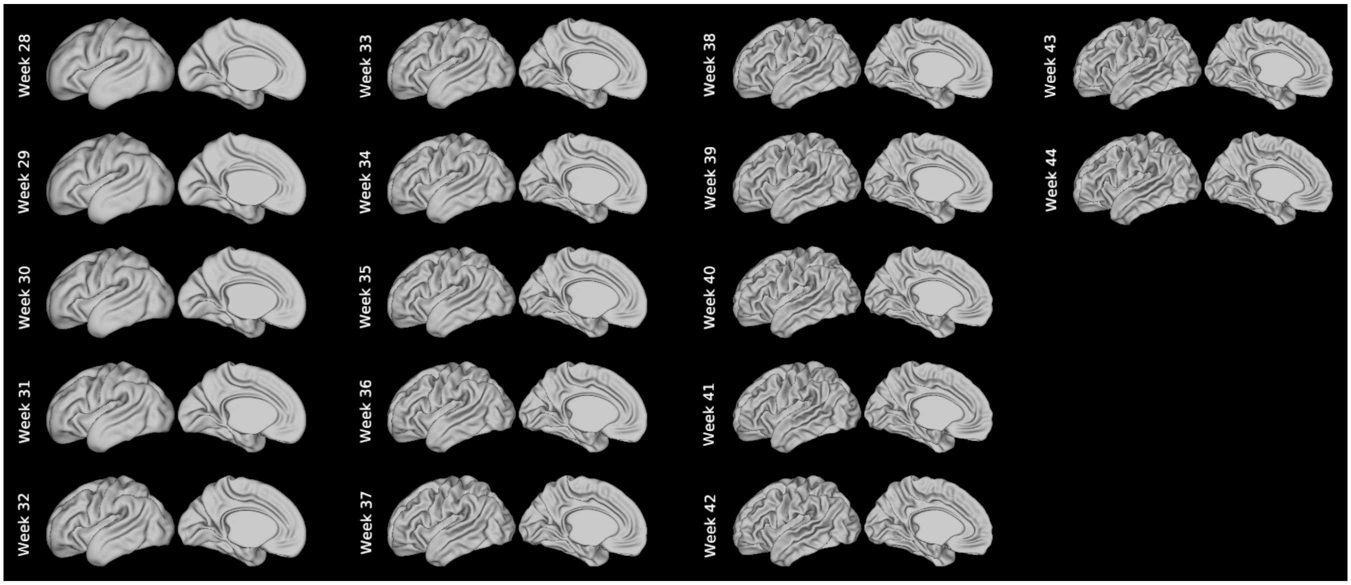


**Fig. S9.** White matter surface for each postmenstrual week of the dhcpSym template. Only the left hemisphere is shown for visualisation purposes. Data at https://balsa.wustl.edu/qx9ZB.

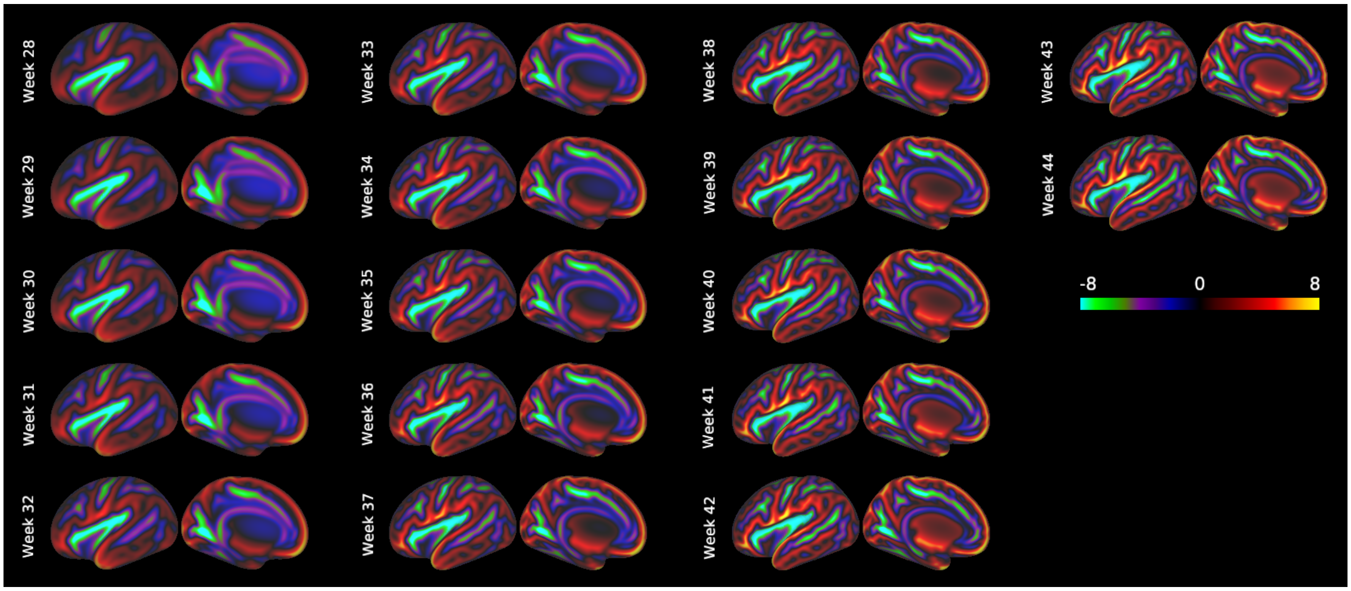


**Fig. S10.** Sulcal depth template for each postmenstrual week of the dhcpSym atlas. Sulcal depth templates are overlaid on a very inflated left hemispheric surface for visualisation purposes. Data at https://balsa.wustl.edu/jNgB2.

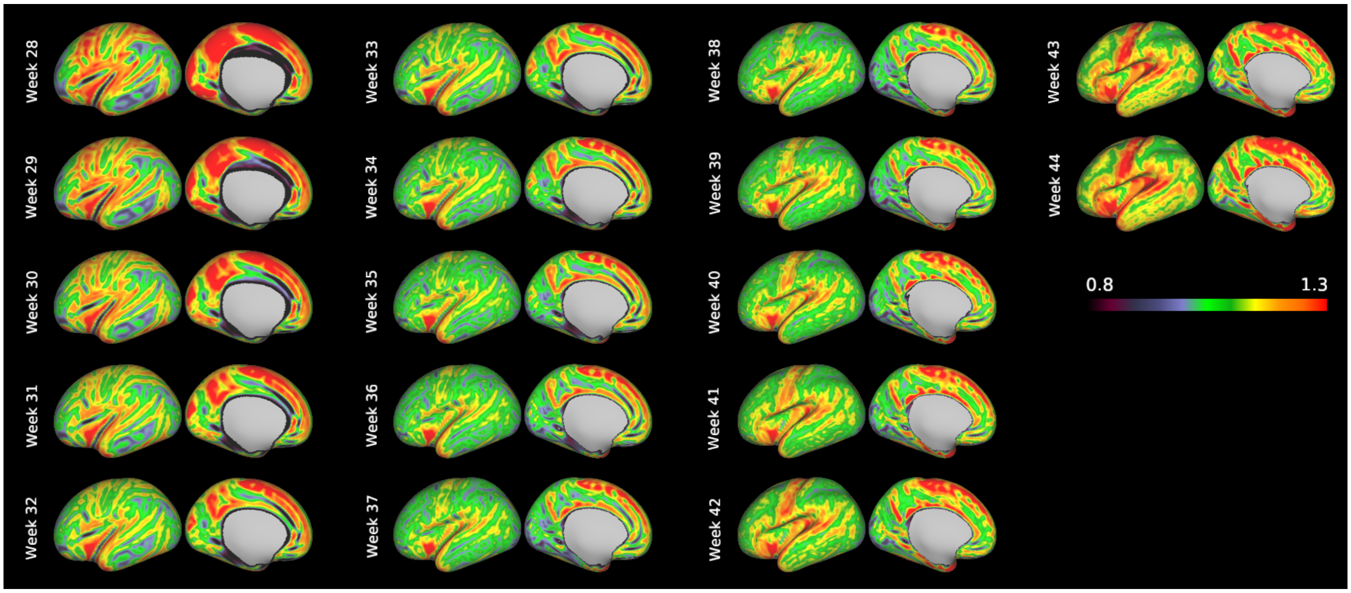


**Fig. S11.** Cortical thickness template for each postmenstrual week of the dhcpSym atlas. Cortical thickness templates are overlaid on a very inflated left hemispheric surface for visualisation purposes. Data at https://balsa.wustl.edu/w8MkL.

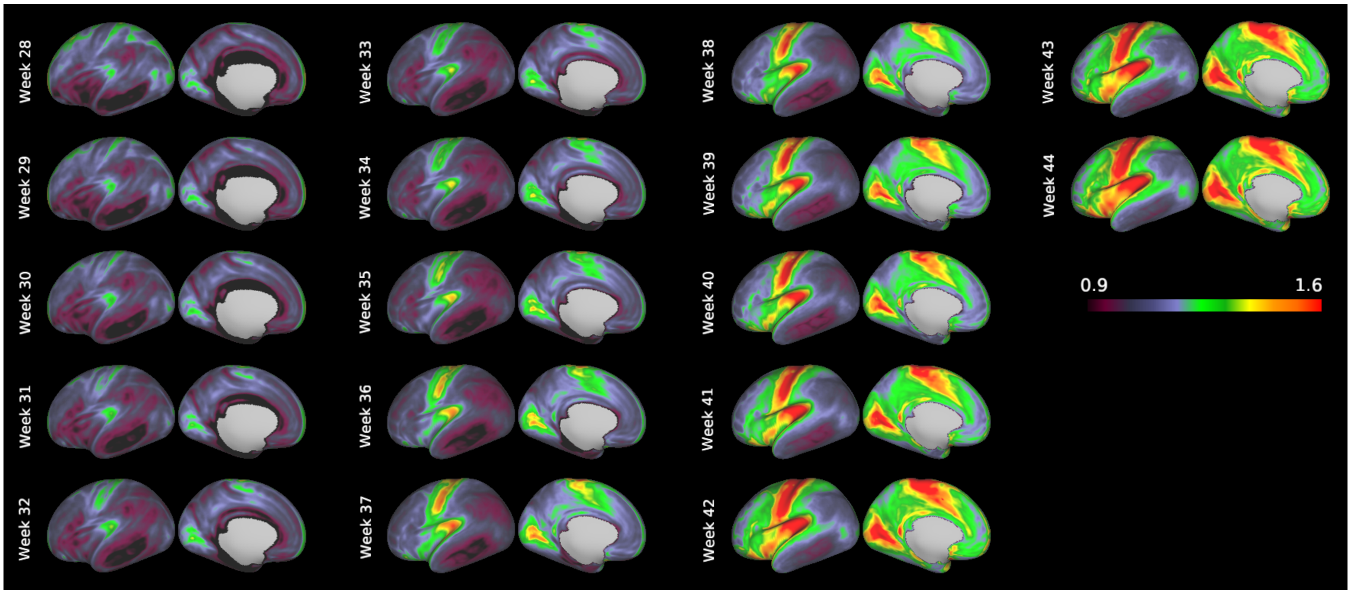


**Fig. S12.** T1w/T2w ratio template for each postmenstrual week of the dhcpSym atlas. T1w/T2w ratio templates are overlaid on a very inflated left hemispheric surface for visualisation purposes. Data at https://balsa.wustl.edu/40Pgj.
